# A non-canonical DNA damage checkpoint response in a major fungal pathogen

**DOI:** 10.1101/2020.08.14.251181

**Authors:** Erika Shor, Rocio Garcia-Rubio, Lucius DeGregorio, David S. Perlin

## Abstract

To protect genome integrity, eukaryotic cells respond to DNA damage by triggering highly conserved checkpoint mechanisms involving the phosphorylation of Rad53/CHK2 kinase. Budding yeast *Candida glabrata*, closely related to model eukaryote *Saccharomyces cerevisiae*, is an opportunistic pathogen characterized by high genetic diversity and rapid emergence of drug resistant mutants. However, the mechanisms enabling this genetic variability are unclear. Here we show that *C. glabrata* cells exposed to DNA damage neither induce CgRad53 phosphorylation nor accumulate in S phase, and exhibit higher lethality than *S. cerevisiae*. Furthermore, time-lapse microscopy showed *C. glabrata* cells continuing to divide in the presence of DNA damage, resulting in mitotic errors and cell death. Finally, RNAseq analysis revealed transcriptional rewiring of the DNA damage response in *C. glabrata* and identified several key protectors of genome stability upregulated by DNA damage in *S. cerevisiae* but downregulated in *C. glabrata*, including PCNA. Together, our results reveal a non-canonical fungal DNA damage response, which may contribute to rapidly generating genetic change and drug resistance.

## INTRODUCTION

DNA damage poses an ever-present threat to living cells. Failure to mount an appropriate response to DNA damage can lead to genetic instability, which has a number of biological and pathological consequences, e.g. contributing to the development of cancer and, in microbial pathogens, affecting the evolution of host-pathogen interplay and the emergence of drug-resistant mutants. The genomes of fungal pathogens in particular have been extensively reported to exhibit genetic instability, particularly during host colonization or under stressful environmental conditions ^1–6^. The types of genetic alterations most commonly described in fungi are aneuploidies and loss of heterozygosity, which occur in diploid or polyploid fungi, such as *Candida albicans* or *Cryptococcus neoformans* ^7^. Although haploid fungi cannot avail themselves of these mechanisms, they can also exhibit extensive genetic variation and rapid emergence of drug resistant mutants ^8–12^, suggesting the existence of other, as yet unknown, mechanisms that enable high genetic “flexibility” in fungal pathogens.

*Candida glabrata* is a haploid budding yeast more closely related to baker’s yeast *Saccharomyces cerevisiae* than to *C. albicans*. Unlike *S. cerevisiae*, however, *C. glabrata* is an obligate human commensal microbe that can become pathogenic and is a leading cause of life-threatening invasive fungal infections in immunocompromised individuals ^13–15^. *C. glabrata* rapidly develops resistance to different antifungal drug classes ^12,16–20^ and is characterized by extremely high genome variation among clinical isolates both in terms of single nucleotide polymorphisms and larger structural variants ^9–12,21^. The documented extensive chromosomal variation among *C. glabrata* clinical isolates resembles the unstable karyotypes and increased gross chromosomal rearrangements observed in *S. cerevisiae* mutants lacking DNA replication checkpoint functions ^22,23^. However, checkpoint activity in *C. glabrata* has not been examined.

Response to DNA damage depends on the cell cycle phase, but damage incurred during the process of DNA replication is considered to be especially detrimental as it can lead to replication fork destabilization, the formation of double-strand breaks at collapsed forks, and inappropriate recombination, resulting in chromosomal rearrangements and cell death ^24,25^. DNA replication checkpoint slows down S-phase progression, stabilizes replication forks, inhibits replication origin firing, and upregulates transcription of DNA repair genes ^26–29^ Cells with unrepaired DNA damage at the end of S-phase also activate the G2/M checkpoint, which arrests cells in mitosis ^30,31^. Although these checkpoints differ with regard to their specific triggers, protein players, and downstream effects, several proteins play key roles in both checkpoints, most notably the Rad53 serine/threonine kinase (CHK2 in higher eukaryotes). Rad53 is a checkpoint effector kinase – upon DNA damage or DNA replication arrest, it is extensively phosphorylated by upstream sensor kinases and by itself ^32,33^, thereupon amplifying the DNA damage signal by phosphorylating dozens of downstream targets ^34–36^. Rad53 phosphorylation is instrumental for virtually all aspects of the DNA damage response ^35,37–41^. Rad53 orthologs are also extensively phosphorylated upon DNA damage in several *non-Saccharomyces* fungal species examined, including *C. albicans, C. neoformans*, and *Schizosaccharomyces pombe* ^42–44^ However, the phosphorylation of Rad53 in *C. glabrata* has not been studied.

In this study we examined the DNA damage response of *C. glabrata*, focusing on Rad53 phosphorylation, cell cycle alterations, and the global transcriptomic response. Interestingly, we did not detect a DNA damage-induced increase in Rad53 phosphorylation in *C. glabrata*. Consistent with this finding, in the presence of DNA damage *C. glabrata* cells did not accumulate in S-phase and proceeded to divide, giving rise to mitotic errors and significant cell death. Finally, using RNAseq we obtained evidence of transcriptional rewiring of the DNA damage response in *C. glabrata*, as well as differential regulation of several key protectors of genome integrity, including proliferating cell nuclear antigen (PCNA). Together, these results reveal previously unappreciated variation in fungal DNA damage responses and have important implications for fungal genome stability, evolution, and emergence of antifungal drug resistance.

## RESULTS

### *C. glabrata* does not induced CgRad53 phosphorylation upon DNA damage

To begin to elucidate the role of the DNA damage checkpoint in *C. glabrata*, we examined the phosphorylation of *C. glabrata* Rad53 (CgRad53; encoded by *CAGL0M02233g*). Rabbit polyclonal antibodies raised against short peptides in the CgRad53 N- and C-termini did not efficiently detect endogenous CgRad53, but adding a plasmid-borne copy of the gene driven by a weak promoter ^45^ resulted in a 4-fold overexpression of *CgRAD53* (quantified by qPCR, data not shown) and robust detection of the protein, allowing us to examine its mobility on SDS-PAGE in the absence and presence of DNA damage. As a control, *S. cerevisiae* Rad53 (ScRad53) was examined as well. Consistent with existing literature, we detected a shift in ScRad53 mobility upon exposure to DNA alkylating agent methyl methanesulfonate (MMS) and oxidative damage by H_2_O_2_ (Figure 1a), reflecting extensive phosphorylation. In contrast, we did not detect a mobility shift for CgRad53, either in MMS or in H_2_O_2_ (Figure 1a). We considered the possibility that CgRad53 phosphorylation occurred rapidly and transiently, so we examined CgRad53 mobility starting at 20 minutes after addition of MMS; however, no mobility shift was detected (Figure 1b). To confirm that *C. glabrata* was experiencing DNA damage in the presence of MMS or H_2_O_2_, we measured the abundance of histone H2A phosphorylated at serine 129 (a.k.a. γH2A.X), a universal marker of DNA damage, particularly double-strand breaks ^46^. We found that γH2A.X was strongly induced both by MMS and by H_2_O_2_ in *C. glabrata* (Figure 1a, 1b). Consistent with reports that *C. glabrata* is highly resistant to oxidative damage ^47^, it required a much higher concentration of H_2_O_2_ than *S. cerevisiae* to cause significant DNA damage (Figure 1a). However, the effects of MMS on γH2A.X levels in *S. cerevisiae* and in *C. glabrata* were similar (Figure 1c). Together, these data indicated that despite efficient induction of DNA damage in *C. glabrata*, CgRad53 mobility did not change, indicating that extensive phosphorylation was not occurring.

**Figure 1.**
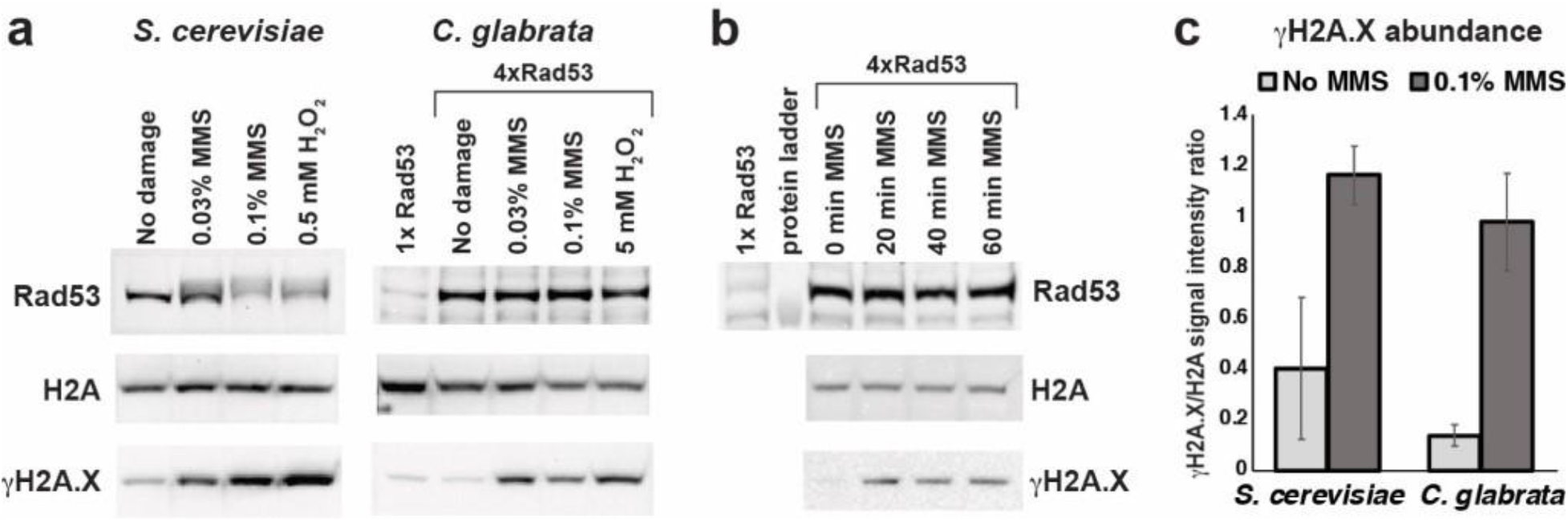
DNA damage induced a change in Rad53 mobility in *S. cerevisiae* but not *C. glabrata*. **a.** Alkylating damage (MMS) and oxidative damage (H_2_O_2_) induced a shift in the mobility of ScRad53 but not CgRad53. Both conditions induced DNA damage in both species, as evidenced by increased abundance of γH2A.X. **b.** MMS induced an increase in γH2A.X abundance by 20 mins post-exposure but did not induce even a transient shift in CgRad53 mobility. **c.** MMS treatment induced DNA damage, as reflected by γH2A.X levels, to similar extent in *S. cerevisiae* and *C. glabrata*. Results were calculated from at least three independent biological replicates for every condition. In **a** and **c** the cells were exposed to the indicated DNA damaging agent for 1 hour.

To further investigate the phosphorylation status of CgRad53 in the absence of presence of DNA damage, we immunoprecipitated endogenous CgRad53 and ScRad53 from untreated and MMS-treated *C. glabrata* and *S. cerevisiae*, respectively, and subjected it to mass spectrometry (MS) analysis (Figure 2a). We identified 346 and 451 unique ScRad53 peptides isolated from untreated and MMS-treated cells, respectively, corresponding to 75% and 82% protein coverage (Supplementary Data 1, Supplementary Figure 1). For CgRad53, we identified 45 and 63 unique peptides obtained from untreated and MMS-treated cells, respectively, corresponding to 44% and 49% protein coverage (Supplementary Data 1, Supplementary Figure 1). Consistent with existing literature, we detected a strong increase in the fraction of ScRad53 phosphopeptides in the MMS-treated sample (Figure 2b, 2c, Supplementary Data). In contrast, and consistent with the Western Blot data (Figure 1a, 1b), the fraction of CgRad53 phosphopeptides did not increase after MMS treatment (Figure 2b, Supplementary Data), supporting the conclusion that *C. glabrata* Rad53 was not significantly phosphorylated upon DNA damage.

**Figure 2.**
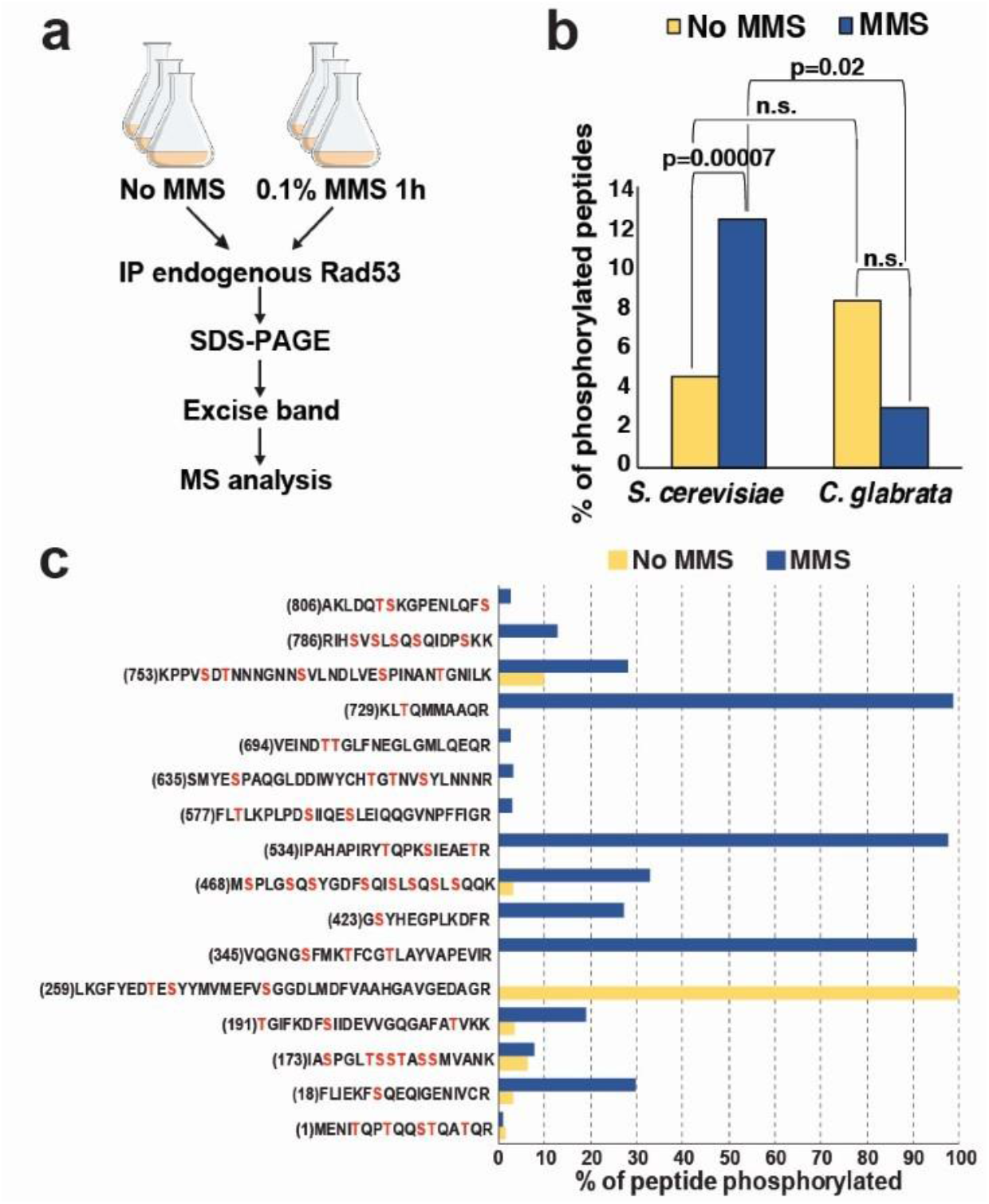
Mass spectrometry (MS) analysis detected extensive DNA damage-induced Rad53 phosphorylation in *S. cerevisiae* but not *C. glabrata*. **a.** Outline of the experiment. **b.** The fraction of phosphorylated Rad53 peptides was significantly increased in *S. cerevisiae* samples, but not *C. glabrata* samples, derived from MMS-treated cells. The p-value was calculated using the chi^2^ test. **c.** Consistent with previous studies, our MS analysis identified extensive DNA damage-induced phosphorylation throughout ScRad53. For each peptide, the total intensity of the phosphorylated forms of that peptide was divided by the total intensity of all forms of that peptide, converted to percentages, and plotted on the Y-axis. The number in parentheses indicates the position of the first residue in the peptide. Serines and threonines are shown in red.

To identify the protein features that may contribute to this lack of phosphorylation, we scrutinized CgRad53 amino acid sequence. CgRad53 is slightly smaller than ScRad53 (767 vs 821 amino acids), but its overall domain organization is similar to ScRad53, containing a kinase domain flanked by two FHA domains (Supplementary Figure 2a). Likewise, the two proteins contain a similar percentage of serines and threonines (Supplementary Figure 2b). However, an examination of the ScRad53-CgRad53 protein alignment revealed that a number of serines and threonines phosphorylated in ScRad53 were not conserved in CgRad53 (Table 1, Supplementary Figure 3). Interestingly, most ScRad53 S/TQ motifs, which are canonical phosphorylation sites for PI-3K-related kinases, such as Mec1 and Tel1, are conserved in CgRad53, with the exception of Thr731 (Table 1). In contrast, all three ScRad53 proline-directed phosphorylation sites (Ser175, Ser375, and Ser774), which are phosphorylated by cyclin-dependent kinases ^48,49^, are not conserved in CgRad53 (Table 1). Likewise, a number of non-canonical (non-ST/Q) Mec1 sites and ScRad53 auto-phosphorylation sites are not conserved in CgRad53 (Table 1). Importantly, the majority of serines and threonines phosphorylated in ScRad53 but lacking conservation in CgRad53 have been shown to be targets of MMS-induced phosphorylation (Table 1). This lack of conservation, together with the results shown above, supports the conclusion that in *C. glabrata* Rad53 is not targeted for extensive DNA damage-induced phosphorylation.

**TABLE 1.**
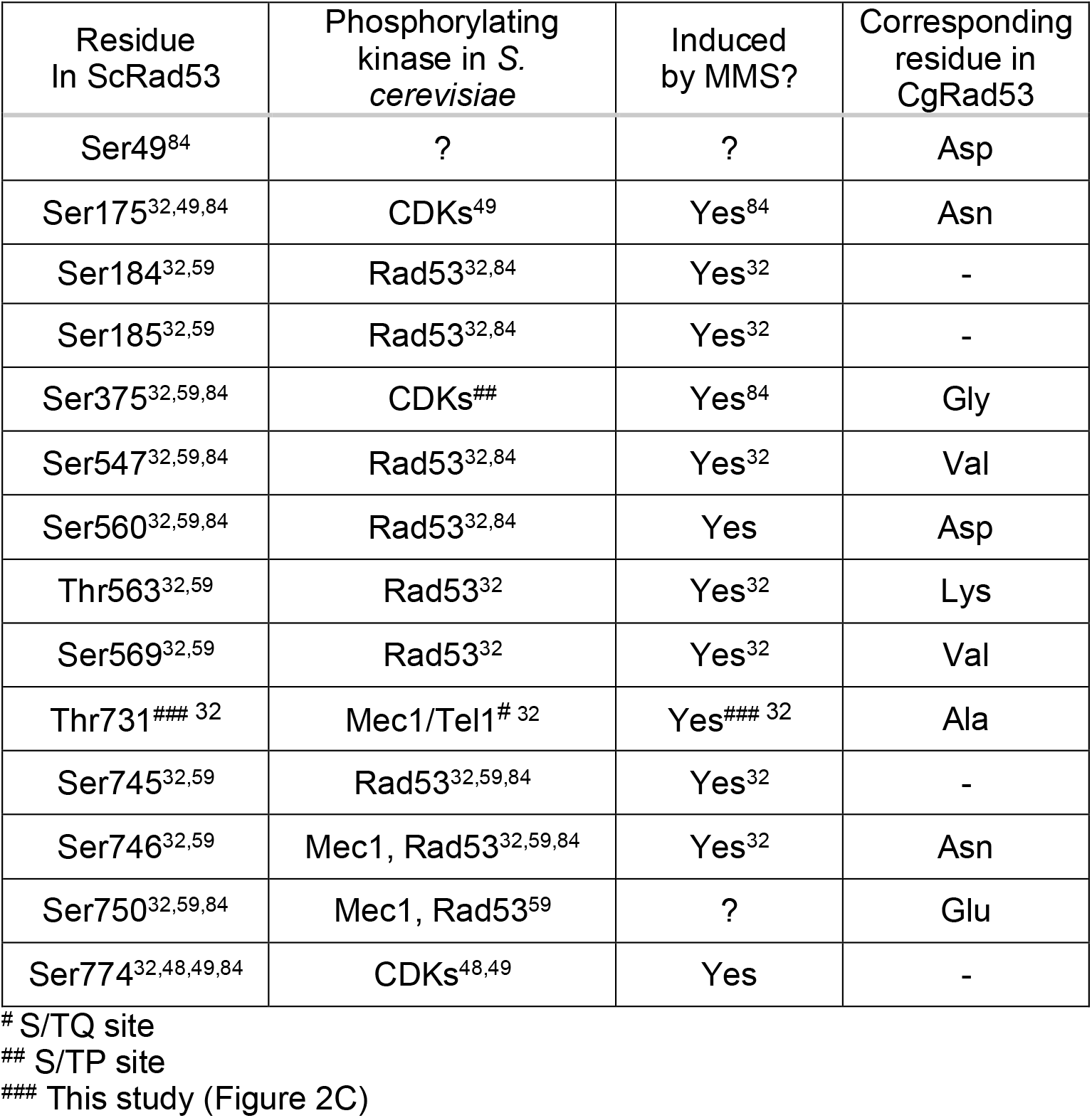
ScRad53 phospho-acceptor sites not conserved in CgRad53

### *C. glabrata* cells do not accumulate in S phase upon DNA damage

A key consequence of DNA damage signaling replication checkpoint activation via Rad53 phosphorylation is the slowing of DNA replication, which allows cells time to repair the damage prior to cell division ^50^. A typical method of detecting this in *S. cerevisiae* involves synchronizing cells in G1 with α-factor and releasing them into DNA damaging conditions. Because *C. glabrata* cells do not arrest in response to mating pheromones ^51^, we used carbon starvation to synchronize *C. glabrata* and *S. cerevisiae* cells in G1 (Figure 3). The synchronized cells were then released into glucose-containing medium in the absence or presence of 0.03% MMS and cell cycle distribution was analyzed for 6 hours by flow cytometry. Hydroxyurea (HU) (100 mM), which inhibits DNA replication by depleting dNTP pools but without inducing DNA damage, was used as a comparator. Consistent with previous reports ^50,52,53^, *S. cerevisiae* cells released into MMS-containing medium significantly slowed down DNA replication, remaining largely accumulated in S-phase by the end of the 6 hours (Figure 3). In contrast, while *C. glabrata* cells were slowed down by the presence of MMS in terms of their entry into S-phase (compare 2h time points for “No drug” and “MMS”, Figure 3), they did not accumulate in S-phase and largely completed DNA replication between 4 and 5 hours after MMS exposure (Figure 3). We did this experiment at both 30°C and 37°C (the optimal *C. glabrata* growth temperature) and obtained identical results (Supplementary Figure 4). Finally, *C. glabrata* released in the presence of HU also delayed the start of DNA replication, but, unlike in the presence of MMS, did not complete it by the end of the 6 hour period, at which point a large proportion of the population still remained in S-phase (Figure 3), whereupon their cell cycle profiles looked similar to HU-exposed *S. cerevisiae* cells (Figure 3). Together, these data show that activation of the S-phase checkpoint by DNA damage (but not by non-damage-associated inhibition of DNA replication) is significantly attenuated in *C. glabrata* compared to *S. cerevisiae*.

**Figure 3.**
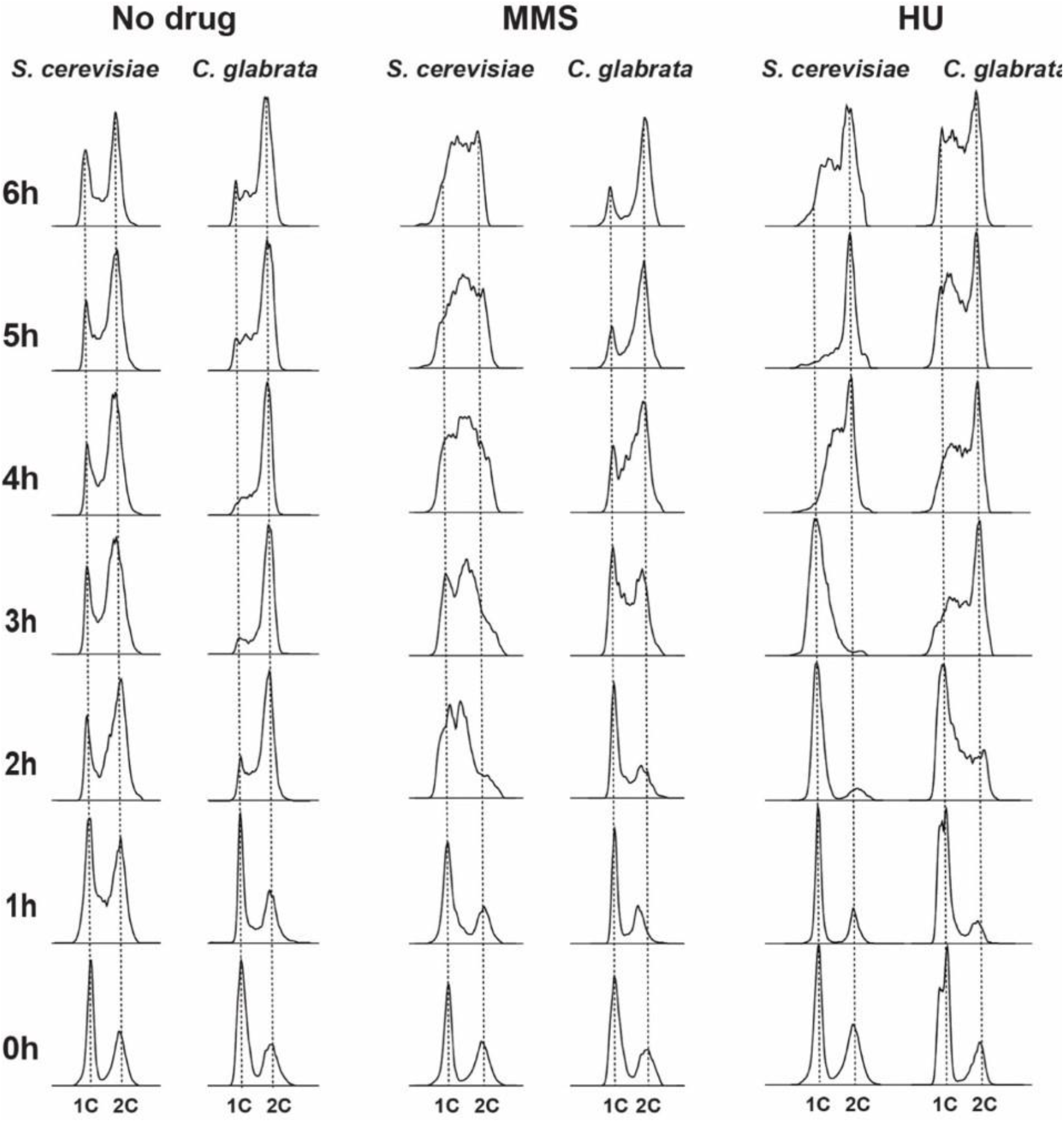
DNA damage induced significant S-phase accumulation in *S. cerevisiae* but not in *C. glabrata*. *S. cerevisiae* and *C. glabrata* cells were synchronized in G1-phase by carbon starvation and released into glucose-containing medium either in the absence or presence of MMS (0.03%) or HU (100 mM). In the presence of MMS, *C. glabrata* completed DNA replication much faster than *S. cerevisiae* cells, which remained accumulated in S-phase by the end of the 6-hour time course. SYTOX Green staining and flow cytometry were used to measure DNA content.

### *C. glabrata* cells undergo aberrant cell divisions and lose viability in response to DNA damage

In *S. cerevisiae*, Rad53-mediated checkpoint signaling is essential for surviving DNA damage, wherein *rad53* mutants (both deletion and point mutants lacking phospho-sites) and other checkpoint mutants proceed with the cell cycle in the presence of DNA damaging agents and exhibit high lethality, presumably due to aberrant replication and division ^53^. Because *C. glabrata* exhibited highly attenuated DNA damage-induced Rad53 phosphorylation and checkpoint activation, we measured its ability to survive DNA damage. We found that whereas at low concentration of MMS (0.01%), viability was moderately and similarly impacted in *S. cerevisiae* and *C. glabrata*, at higher MMS concentrations (0.03% and 0.1%), *C. glabrata* was significantly more sensitive than *S. cerevisiae*, exhibiting several orders of magnitude higher lethality after 8 hours in 0.1% MMS (Figure 4a).

**Figure 4.**
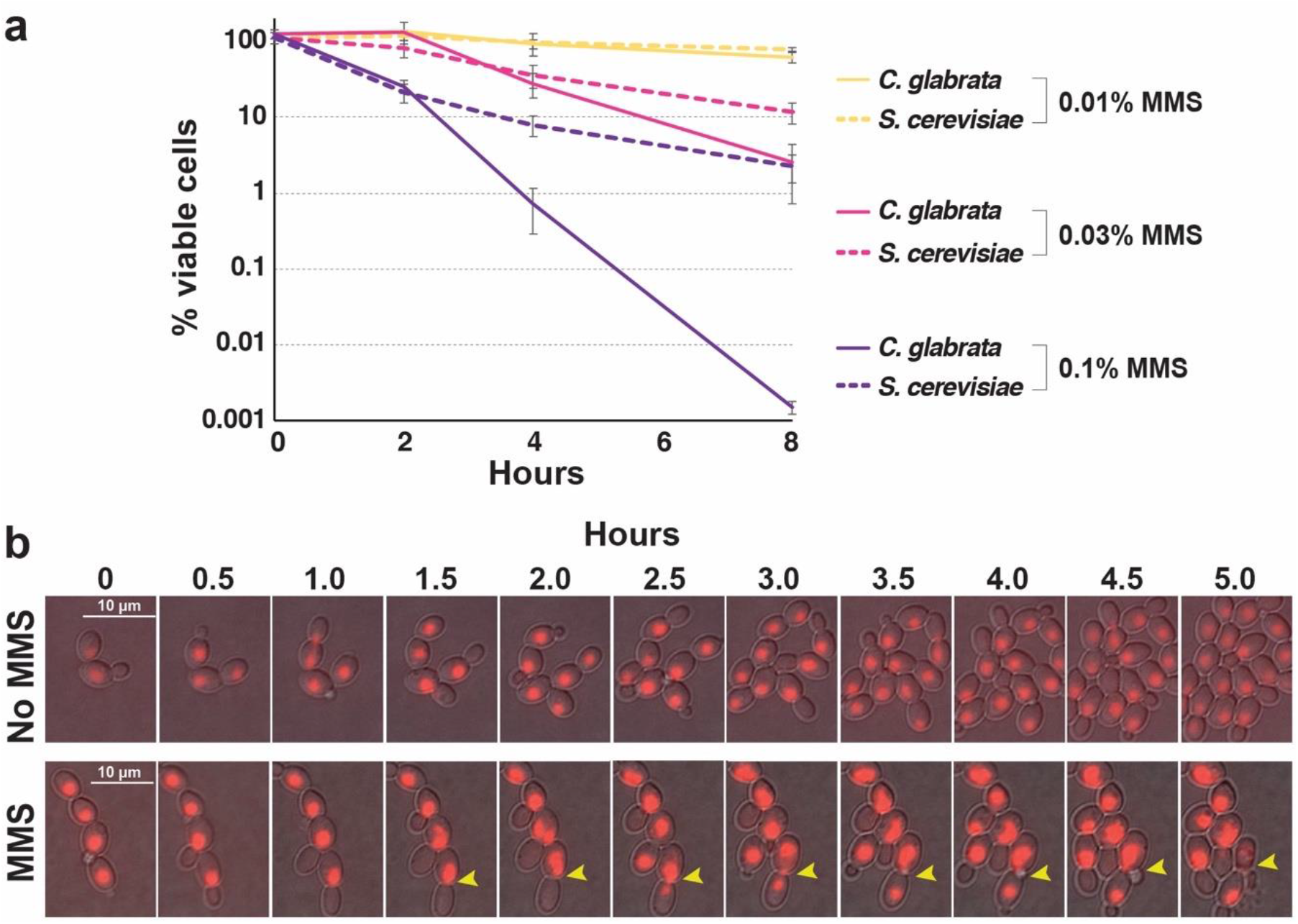
*C. glabrata* exhibited high lethality and aberrant mitoses in the presence of DNA damage. **a.** *C. glabrata* cells are more sensitive to high levels of DNA damage than *S. cerevisiae* cells. Cells were cultured in the presence of indicated concentrations of MMS, harvested at the indicated time points, counted, and plated on drug-free YPD plates. Viability counts were obtained by dividing the number of resulting colonies by the number of plated cells. Results were calculated from at least three biological replicates for every time point. **b.** Time-lapse microscopy detected *C. glabrata* cells dividing in the presence of 0.03% MMS, including aberrant nuclear divisions. The cells, carrying an NLS-RFP construct to fluorescently mark nuclei, were pipetted onto YPD-agarose pads, sealed, and imaged for 6 hours at 10-minute intervals. 30-minute intervals are shown. Yellow arrowheads indicate a cell where nuclear material was unequally distributed into mother and daughter cells. Both mother and daughter cells subsequently budded, but the mother cell burst. These and other groups of cells are shown in Supplementary Movies 1-5.

To gain insight into the causes of lethality in MMS-treated *C. glabrata* cells, we used time-lapse microscopy to track cell division of *C. glabrata* cells with fluorescently marked nuclei (NLS-RFP; Supplementary Videos 1-5). We observed that whereas the presence of 0.03% MMS significantly slowed down the rate at which new buds emerged, nevertheless, 2-3 hours after the addition of MMS a number of cells proceeded with nuclear division and mitosis (Figure 4b, Supplementary Videos 1-5). Furthermore, we were able to observe aberrant mitoses wherein nuclear content was distributed unequally between mother and daughter cells prior to cytokinesis (Figure 4b, yellow arrowhead). Despite this unequal distribution of nuclear content, both mother and daughter cells proceeded to bud; however, the mother cell subsequently “exploded” (Figure 4b, yellow arrowhead). These observations, together with the cell cycle distribution analysis (Figure 3) and cell viability measurements (Figure 4a), show that *C. glabrata* cells do not significantly activate the DNA damage checkpoint, that many of them proceed with S-phase and cell division even in the presence of DNA damage, and as a result lose viability due to aberrant mitoses.

### A rewiring of the transcriptional response to DNA damage in *C. glabrata*

A key part of the cellular response to DNA damage is activated Rad53 phosphorylating multiple transcription factors, which in turn alter the expression of hundreds of genes, e.g. downregulating genes involved in growth and cell cycle progression and upregulating genes involved in stress responses and DNA repair^35,40^. To ask whether a similar transcriptional response exists in *C. glabrata*, we cultured both *C. glabrata* and *S. cerevisiae* in the absence or presence of 0.1% MMS for one hour, isolated total RNA, and analyzed it by RNAseq. As reported previously, over 2000 genes were up- or down-regulated by DNA damage in *S. cerevisiae* (Figure 5a, Supplementary Data 2), and these transcriptional changes were consistent with those published previously (Supplementary Figure 5) ^35,40^. Over 2000 genes were also up- and downregulated by DNA damage in *C. glabrata* (Figure 5a) and, interestingly, there was a high degree of concordance between the expression changes of orthologous genes present in both species (4797 genes; Figure 5a, 5b). This concordance was especially strong for genes downregulated by MMS: in both *S. cerevisiae* and *C. glabrata* these genes were strongly enriched for those involved in protein synthesis, e.g. translation, ribosome biogenesis, and rRNA processing (Supplementary Figure 6). This downregulation of pro-growth genes was consistent with previous reports ^35,40^ and with our conclusion that in the presence of MMS *C. glabrata* was experiencing DNA damage-induced stress. Interestingly, gene categories induced by MMS were more diverse in *C. glabrata* than in *S. cerevisiae*. Both species induced genes involved in protein degradation and stress responses; however, *C. glabrata* also induced orthologs of genes, which in *S. cerevisiae* are involved in sporulation and meiosis (Supplementary Figure 6). This observation was intriguing and unexpected because mating and meiosis have not been detected in *C. glabrata* to date.

**Figure 5.**
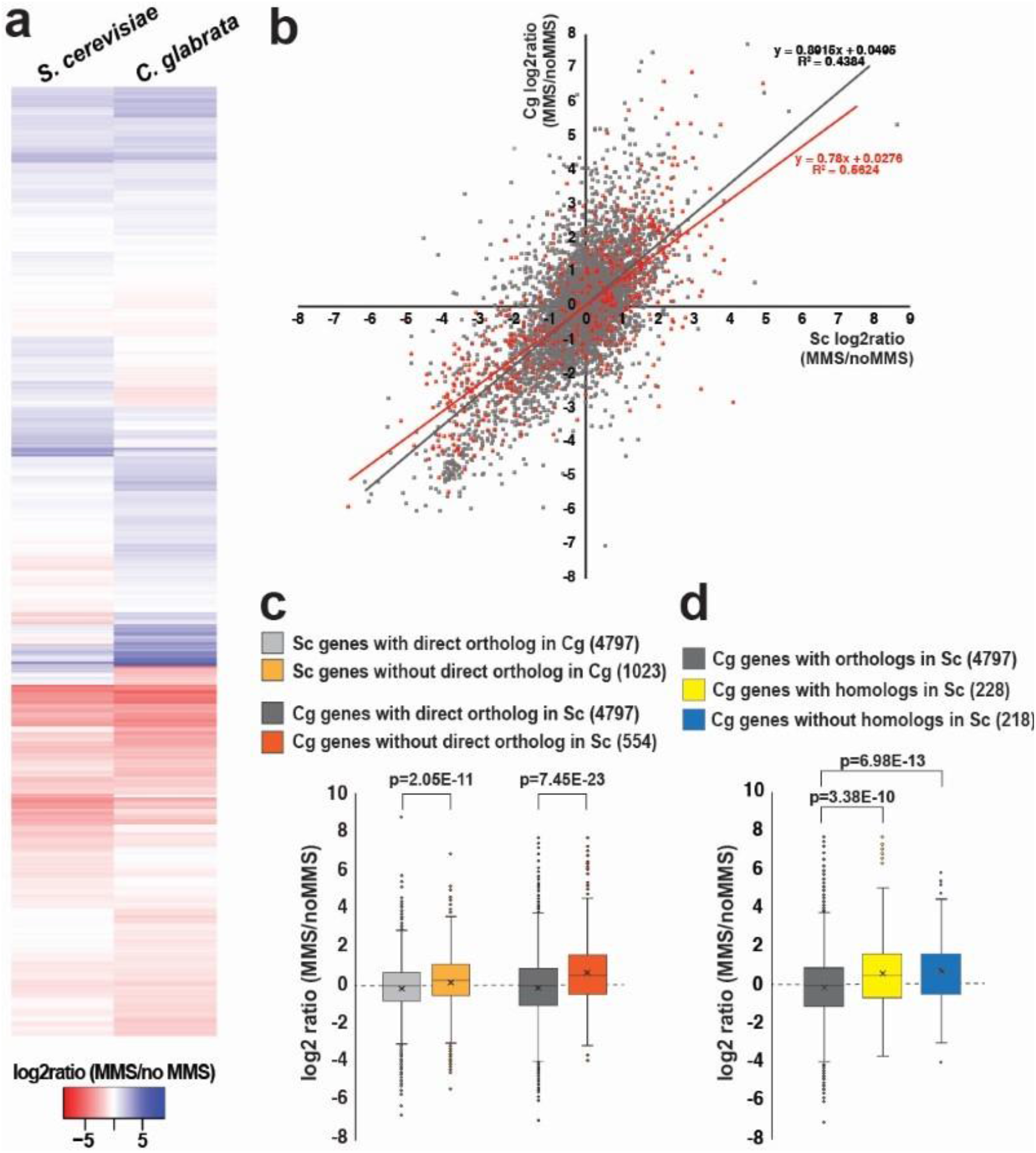
RNAseq revealed evidence of transcriptional rewiring of the DNA damage response in *C. glabrata* relative to *S. cerevisiae*. *C. glabrata* and *S. cerevisiae* cells were treated with 0.1% MMS for 1 hour, then total RNA was isolated from both untreated and treated cells and subjected to RNAseq analysis. Three biological replicates of every condition were analyzed, with the exception of “*S. cerevisiae* no MMS”, for which one of the samples had poor RNA quality and was not processed further. **a.** A heatmap of the “MMS/no MMS” log2 ratios for *S. cerevisiae* genes and their *C. glabrata* orthologs. **b.** A scatterplot where each gene is represented by a dot and its “MMS/no MMS” log2 ratio for *S. cerevisiae* is plotted on the x-axis and for *C. glabrata* on the y-axis. Genes whose expression is regulated by Rad53 in *S. cerevisiae* are indicated in red. **c.** In both *C. glabrata* and *S. cerevisiae* genes that lack a direct ortholog in the other species are induced more strongly by DNA damage. **d.** Both *C. glabrata* genes that have homologs but not direct orthologs in *S. cerevisiae* and *C. glabrata* genes that have no homologs in *S. cerevisiae* tend to be upregulated by MMS. The p-values were calculated by an unpaired two-tailed t-test. The *S. cerevisiae-C. glabrata* ortholog list was downloaded from http://www.candidagenome.org/download/homology/orthologs. In **c** and **d** the number in parentheses indicates the number of genes in the corresponding category.

We also specifically examined genes whose DNA damage-induced expression changes (either up- or downregulation) in *S. cerevisiae* are known to be Rad53-dependent ^35^. Interestingly, the majority of these genes’ orthologs were also responsive to DNA damage in *C. glabrata*, and their overall response to MMS was similar in *C. glabrata* to that in *S. cerevisiae* (Figure 5b, red dots). This result suggested that this set of genes had undergone transcriptional “rewiring” in *C. glabrata*, whereby their transcription was robustly induced or repressed by DNA damage within the timeframe (one hour) where CgRad53 phosphorylation was not induced, and that therefore these changes may have been mediated by factors other than Rad53.

We also individually examined several canonical Rad53 target genes, i.e. those whose transcription has long been known to be strongly induced by DNA damage in Rad53-dependent manner, specifically ribonucleotide reductase subunit *RNR3* and ribonucleotide reductase inhibitor *HUG1* ^54,55^. However, we found that *C. glabrata* genome did not contain direct orthologs of either *ScRNR3* or *ScHUG1*. This prompted us to compare DNA damage-induced transcriptional changes of genes that had a direct ortholog in the other species (~73% of all *S. cerevisiae* genes) to those that lacked such orthologs. The *S. cerevisiae-C. glabrata* ortholog information was obtained from the *Saccharomyces* Genome Database (www.yeastgenome.org) and the *Candida* Genome Database (www.candidagenome.org). Interestingly, we found that both in *S. cerevisiae* and *C. glabrata* genes that lacked a direct ortholog in the other species were significantly more likely to be induced by DNA damage than genes that did have an ortholog (Figure 5c). To probe this phenomenon further, we focused on the genes in *C. glabrata*, which, according to www.candidagenome.org, did not have a direct ortholog in *S. cerevisiae*. These genes generally could be subdivided into two categories: those that had homologs in *S. cerevisiae* and those, for which BLAST searches had revealed no homologs in *S. cerevisiae* (Supplementary Data 2). Interestingly, we found that both groups tended to be significantly more induced by MMS than genes with direct orthologs in *S. cerevisiae* (Figure 5d). In particular, we identified 28 *C. glabrata* genes lacking identifiable *S. cerevisiae* homologs that were at least two-fold downregulated by MMS, and 77 such genes that were at least two-fold upregulated by MMS (Supplementary Data 2). This result suggested that the transcriptional response to DNA damage has been diverging in *S. cerevisiae* and *C. glabrata* during evolution, consistent with other evidence of transcriptional rewiring.

### DNA damage differentially regulates the expression of proliferating cell nuclear antigen (PCNA) in *C. glabrata* and *S. cerevisiae*

As is evident from Figure 5a, a number of genes were differentially regulated in *S. cerevisiae* and *C. glabrata*. We defined “differential regulation” as a difference of at least 2 logs, or 4-fold, in expression change. For instance, by this criterion, a gene whose expression was unchanged by MMS in *S. cerevisiae* would be considered differentially regulated in *C. glabrata* if its expression was induced or repressed at least 4-fold in that organism. Genes that were upregulated by MMS in *C. glabrata* relative to *S. cerevisiae* were enriched for sulfate assimilation (likely in response to MMS), certain types of amino acid metabolism, and meiosis, whereas genes that were downregulated by MMS in *C. glabrata* relative to *S*. cerevisiae were enriched for nucleotide/nucleoside metabolism (Supplementary Figure 7).

We were particularly interested in differentially regulated genes involved in DNA metabolism and genome stability and identified 17 such genes that were up-regulated in *C. glabrata* relative to *S. cerevisiae* and 22 such genes that were down-regulated in *C. glabrata* relative to *S. cerevisiae* (Figure 6a). Interestingly, the latter set contained a number of genes involved in the initiation and progression of DNA replication, including PCNA *(POL30)*, a subunit of the pre-replicative complex *(CDC6)*, several subunits of the MCM replicative helicase, a subunit of DNA polymerase delta *(POL31)*, and Okazaki fragment processing exonuclease *(RAD27)*. Several of these factors also play key roles in maintaining the stability of DNA replication forks in the presence of DNA damage, most notably PCNA, which mediates multiple interactions between the replisome and various DNA repair complexes ^56,57^. *POL30* transcript abundance was induced over 2-fold by MMS in *S. cerevisiae*, consistent with other studies ^35,40^, but repressed by over 8-fold in *C. glabrata* (Supplementary Data 2). Because a decrease in PCNA abundance is expected to drastically affect the stability of DNA the replisome, especially in the presence of DNA damage, we sought to confirm that PCNA expression was affected not only at mRNA but also at protein level. Indeed, we found that upon MMS treatment PCNA abundance increased in *S. cerevisiae* but decreased in *C. glabrata*, consistent with RNAseq results (Figure 6b, 6c). Together, these results showed that several genes with key roles in maintaining replication fork integrity, including PCNA, are upregulated in *S. cerevisiae* but downregulated in *C. glabrata* in response to DNA damage, with likely profound implications on genome stability.

**Figure 6.**
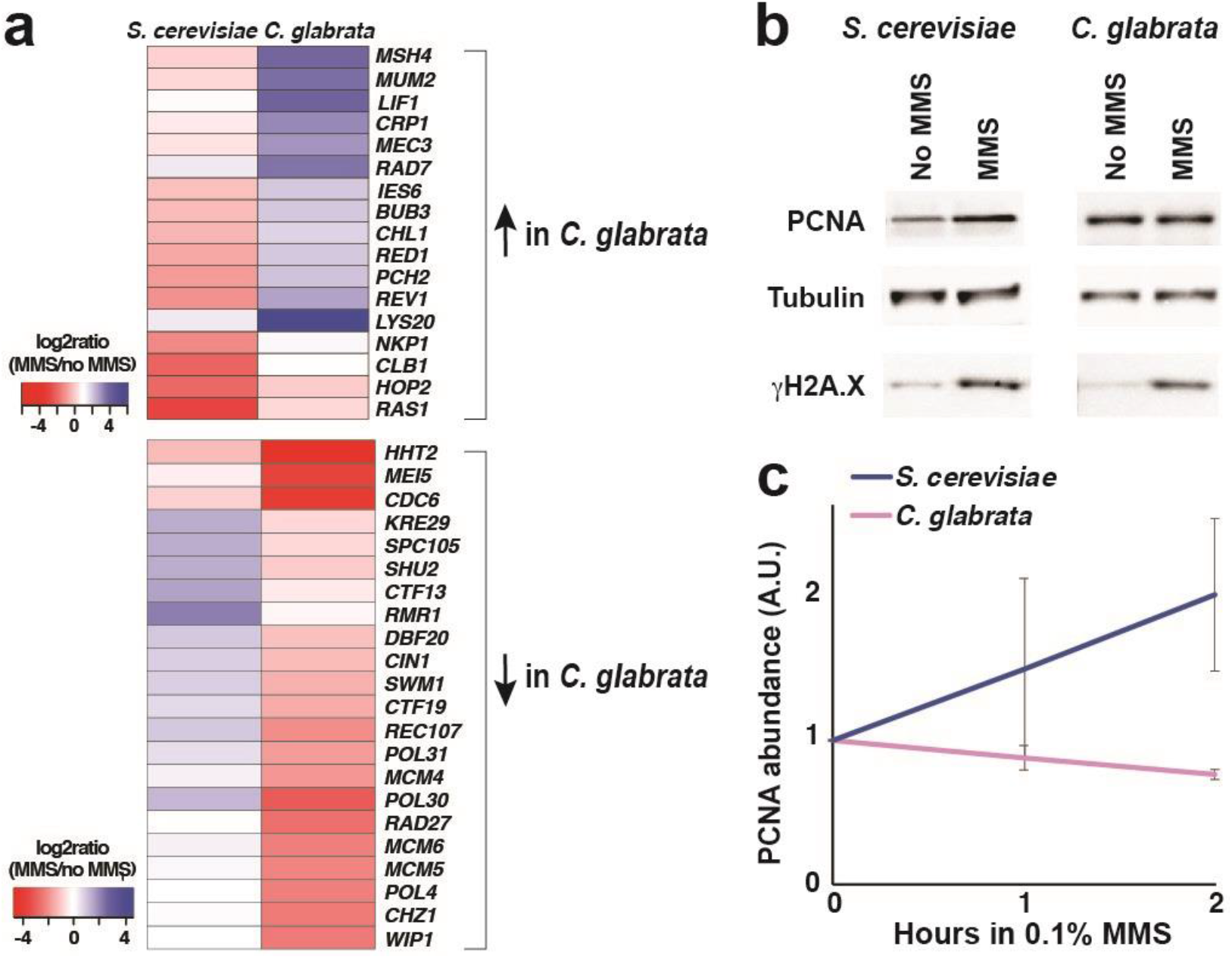
Expression of PCNA is upregulated by DNA damage in *S. cerevisiae* but downregulated in *C. glabrata*. **a.** Heatmaps of genes involved in maintenance of genome stability and differentially regulated by DNA damage in *S. cerevisiae* and *C. glabrata*. **b.** PCNA protein levels increase in response to DNA damage in *S. cerevisiae* but not in *C. glabrata*. Cells were treated by 0.1% MMS by 2 hours, then harvested for total cell lysates and Western blotting. Tub = alpha-tubulin. **c.** Quantification of Western blot data from at least three biological replicates (Image J). For every condition, PCNA abundance was normalized to that of alpha-tubulin.

## DISCUSSION

Our study presents the first examination of the DNA damage checkpoint in the opportunistic fungal pathogen *C. glabrata*. Although *C. glabrata* is closely related to *S. cerevisiae*, we found a number of important differences between the two organisms’ DNA damage responses. Unlike *S. cerevisiae, C. glabrata* did not induce Rad53 phosphorylation or accumulate in S-phase upon DNA damage, indicating reduced activation of DNA replication checkpoints. Consistent with attenuated checkpoint signaling, *C. glabrata* exhibited higher lethality in the presence of DNA damage, and time-lapse microscopy detected evidence of aberrant mitoses under these conditions. Finally, we obtained evidence of diverged transcriptional responses to DNA damage in *C. glabrata* and *S. cerevisiae*, including differential regulation of some key protectors of replication integrity, such as PCNA. Together, these results reveal a new variation in eukaryotic DNA damage responses and expand our understanding of factors influencing fungal genetic stability, evolution, and emergence of antifungal drug resistance.

Mechanistic studies in *S. cerevisiae* have shown that upon DNA damage or replication fork stalling Rad53 is recruited by adaptor proteins (Rad9 or Mrc1) to activated DNA damage sensor kinases (Mec1/Tel1), which phosphorylate Rad53 at both canonical (S/TQ) and non-canonical sites ^32,58,59^. According to current models, phosphorylated Rad53 accumulates at the sites of DNA damage, further extensively auto-phosphorylates in trans, and then diffuses away to phosphorylate multiple downstream targets ^33,35,36,60–63^. Interestingly, we found most S/TQ sites are conserved between ScRad53 and CgRad53, suggesting that Mec1/Tel1 phosphorylation of CgRad53 probably occurs and plays an important role. Another piece of evidence suggesting that Mec1 is likely active in *C. glabrata* is the observed extensive DNA damage-induced phosphorylation of histone H2A-Ser129 (γH2A.X), which is phosphorylated by Mec1 at the sites of damage ^64,65^. In contrast, a number of ScRad53 autophosphorylation sites are not conserved in CgRad53. Thus, it is possible that the initial Mec1-catalyzed phosphorylation of Rad53 takes place in *C. glabrata*, but that it does not lead to the same type of autophosphorylation and activation of this effector kinase and consequently does not trigger the same degree of checkpoint activation.

We observed DNA damage-triggered induction or repression of most *C. glabrata* genes whose *S. cerevisiae* orthologs are dependent on Rad53, suggesting that these *C. glabrata* genes are still under checkpoint control. However, a lack of CgRad53 DNA damage-induced phosphorylation suggests that in *C. glabrata* these genes may not be regulated by Rad53. Indeed, Rad53/CHK2 is not the only DNA damage checkpoint effector kinase in eukaryotic cells. Chk1 (CHK1 in higher eukaryotes) is another serine/threonine effector kinase, which, although playing a minor role in *S. cerevisiae*, has an important role in the DNA damage and replication checkpoint responses of higher eukaryotes and *S. pombe* ^66^. Another effector kinase expressed by yeast cells is Mek1 (meiotic effector kinase), which in *S. cerevisiae* is meiosis-specific and involved in sensing the number of double-strand breaks (DSBs), channeling their repair to promote the appropriate level and distribution of crossovers between homologous chromosomes, and delaying entry into meiosis I until DSB repair has been completed ^67^. Interestingly, and possibly relatedly, we have detected an upregulation of meiosis and sporulation genes upon DNA damage in *C. glabrata*. This observation is intriguing because mating and sporulation have not been detected in *C. glabrata*, although genomic studies suggest that they do happen, albeit extremely rarely ^9,68^. Also, interestingly, both CHK1 and MEK1 are transcriptionally upregulated more strongly in *C. glabrata* than in *S*. cerevisiae by DNA damage, whereas *RAD53* is similarly and very moderately upregulated in both (Supplementary Data 2). Any possible roles of Chk1 and Mek1 effector kinases in the DNA damage response of *C. glabrata* will be elucidated in further studies.

In *S. cerevisiae* Rad53 is phosphorylated not only by DNA damage sensor kinases Mec1 and Tel1, but also by two cell cycle regulators: cyclin-dependent kinase Cdc28/Cdk1 and Polo-like kinase Cdc5, which phosphorylate ScRad53 at three proline-directed sites (Ser175, Ser375, and Ser774) ^48,49^. Interestingly, we found that none of these three phospho-acceptor amino acids are conserved in CgRad53. This lack of conservation is difficult to interpret at present, however, because the role of this phosphorylation in *S. cerevisiae* is still unclear. On the one hand, alanine substitution mutations at these sites do not cause defects in DNA damage-induced Rad53 phosphorylation or DNA damage sensitivity ^48,49^; on the other, phosphorylation of all three of these residues is induced by MMS ^32,59^. These alanine substitutions have a few reported phenotypes, including accelerated cellular recovery from a persistent DNA damage checkpoint signal ^49^ and defects in cell wall integrity ^48^. The latter may be important in *C. glabrata*, as its cell wall is the principal mediator of its interaction with the host and a target of antifungal drugs ^69,70^. Interestingly, a role of checkpoint proteins in morphogenesis and cell wall integrity has been reported in *S. cerevisiae* ^71^. Thus, it will be of interest to examine the role these factors play in cell wall maintenance, drug resistance, and virulence in *C. glabrata*.

Our transcriptome analysis identified a number of genes involved in maintaining genome stability differentially regulated in *S. cerevisiae* and *C. glabrata*. In this study we focused on PCNA for a number of reasons. First, its transcription was strongly differentially affected by MMS, being induced by over two-fold in *S. cerevisiae* and repressed by over eight-fold in *C. glabrata*. The induction of PCNA expression by DNA damage in *S. cerevisiae*, which has been reported before and shown to be dependent on ScRad53 ^35^, is not surprising. Whereas originally defined as the processivity factor for DNA polymerases, PCNA is now known to regulate virtually every aspect of chromosomal maintenance, including DNA replication, recombination, repair, and chromatin structure (reviewed in ^56,57^). *POL30* is an essential gene in *S. cerevisiae*, but a number of mutant alleles have been generated and shown to exhibit aberrant DNA damage repair and elevated rates of mutation and recombination, among other defects ^72–75^ A “Decreased Abundance by mRNA Perturbation” (DAmP) *POL30* allele has also been generated and, while its phenotype with respect to genome stability has not been described, large-scale genetic analyses suggest that it behaves similarly to null mutants in non-essential DNA replication genes, such as *RAD27, POL32*, and *ELG1* ^76^. Further studies are necessary to understand why *C. glabrata* suppresses PCNA expression at a time when it appears to be especially critical for repair of DNA damage and preventing mutagenesis and genomic instability.

Both pathogenic and nonpathogenic fungi are characterized by extensive genetic diversity and ability to adapt to new environments ^77,78^. In fungal species that can associate with humans, this adaptability is important for microevolution within the host and can translate into the development of drug-resistant infections ^3,6,79,80^. In diploid or polyploid fungi, such as *C. albicans* and *C. neoformans*, environmental stress associated with passage through a mammalian host or antifungal drug exposure leads to increased genetic instability, most notably aneuploidies and loss of heterozygosity (LOH) ^1,2,5^. Because *C. glabrata* is haploid, it cannot avail itself of these mechanisms. Furthermore, *C. glabrata* appears to propagate almost exclusively clonally, so it also cannot use meiotic recombination to promote genetic diversity. Yet, *C. glabrata* genome analyses indicate the occurrence of frequent chromosomal rearrangements ^9–11,21^, and our study suggests that these rearrangements may be facilitated by a “lax” DNA damage checkpoint mechanism. *C. glabrata* is the first obligate haploid commensal/pathogenic fungus whose checkpoint activity has been examined. Thus, it will be of interest to examine whether other haploid fungi, for example *C. auris*, which is likewise characterized by extensive genetic variability and high prevalence of antifungal drug resistance ^8^, may also have a non-canonical DNA damage checkpoint. Finally, although such non-canonical checkpoint mechanisms may facilitate genome instability and emergence of drug-resistant strains, they may also present an exploitable therapeutic opportunity to selectively target checkpoint-deficient cells. For instance, strategies are being evaluated for treating checkpoint-deficient human cancers where it may be possible to inhibit CHK1 in CHK2-deficient cancers or vice versa ^81^. It would be of interest to investigate similar approaches in fungi, especially those with non-canonical checkpoint responses.

## MATERIALS AND METHODS

### Yeast strain growth and handling

Common *C. glabrata* reference strain ATCC2001 (also known as CBS138) and *S. cerevisiae* strain W4069-4C (MAT**a**, W303 genetic background, gift of the Rothstein lab) were used for all experiments. Cells were cultured in standard rich medium (YPD). *S. cerevisiae* cells were grown at 30°C and *C. glabrata* cells were grown at 37°C, which are optimal growth temperatures for these organisms. To rule out the effects of temperature on cell cycle progression in the presence of DNA damage, we performed this experiment with *C. glabrata* both at 37°C and 30°C and observed no differences (Figure 3, Supplementary Figure 4).

### Western blotting

Whole cell lysates were prepared by trichloroacetic acid (TCA) precipitation. Briefly, cell pellets were resuspended in 20% TCA, broken by bead beating, washed twice with 5% TCA, and then proteins were pelleted and resuspended in SDS-PAGE loading buffer. Samples were incubated at 95°C for 5 mins and centrifuged prior to loading on acrylamide gels. 8% gels were used to detect Rad53 and 12% gels were used to detect histone H2A, α-tubulin, and PCNA. The following antibodies were obtained commercially: anti-ScRad53 (Abcam ab150018), anti-H2A (Active Motif #39945), anti-γH2A.X (Abcam ab15083), anti-PCNA (Abcam ab221196), and anti-α-tubulin (ThermoFisher Scientific MA1-80189). Rabbit antibodies against an N-terminal peptide (IPIKDMEVDVEQIA) and a C-terminal peptide (GIPNEERSVTSQTE) of CgRad53 were raised by Genscript (https://www.genscript.com). To help detect CgRad53 by Western blot, *C. glabrata RAD53* ORF was subcloned into plasmid pCN-EGD2 (obtained from Addgene) downstream of the weak *EGD2* promoter ^45^. Cells carrying the resulting plasmid were processed for Western blotting as described above.

### Rad53 phosphorylation analysis by MS

Rad53 was immunoprecipitated from *S. cerevisiae* and *C. glabrata* total cell lysates using the anti-Rad53 antibodies described in the previous section and Protein A magnetic beads (New England Biolabs). The immunoprecipitated samples were run on 8% acrylamide gels, stained with GelCode Blue reagent (ThermoFisher Scientific), whereupon an area of roughly 1 cm x 1 cm around the Rad53 band was excised and sent for MS analysis at the Georgetown University Proteomics Shared Resource facility (https://lombardi.georgetown.edu/research/sharedresources/pmsr/proteomics), where the samples were destained and subjected to in-gel proteolytic digestion with trypsin/Lys-C mixture (Promega). The digests were extracted, analyzed by nanoUPLC-MS/MS using the TripleTOF 6600 mass spectrometer, and mass spectra were recorded with Analyst TF 1.7 software. Data files were submitted for simultaneous searches using Protein Pilot version 5.0 software (Sciex) utilizing the Paragon and Progroup algorithms and the integrated false discovery rate (FDR) analysis function. MS/MS data were searched against either ScRad53 or CgRad53 protein databases, as appropriate. Phosphorylation emphasis was chosen as a special factor. The proteins were inferred based on the ProGroupTM algorithm associated with the ProteinPilot software. The detected protein threshold in the software was set to the value which corresponded to 1% FDR. All peptides were filtered with confidence to 5% FDR, with the confidence of phosphorylation sites automatically calculated. The Georgetown University Proteomics Shared Resource facility then provided a list of recovered peptides, their intensities, and their post-translational modifications to the investigators, who used it calculate the relative abundance of phosphorylated peptides in every sample.

### Cell cycle analysis

To synchronize *S. cerevisiae* and *C. glabrata* in the G1-phase of the cell cycle, exponentially growing YPD cultures were shifted to YP medium (no dextrose) and cultured for 18 hours. At that point, cells were resuspended in YPD in the absence or presence of 0.03% MMS or 100 mM HU and cultured for another 6 hours. Aliquots were fixed with 70% ethanol at every hour. Prior to analysis by flow cytometry, the samples were pelleted and resuspended in PBS, sonicated, and treated with an RNase cocktail (Fisher Scientific). The cell counts in each sample were measured and adjusted to the same cell concentration, followed by addition of SYTOX Green (ThermoFisher Scientific) and flow cytometric analysis using the BD Fortessa instrument.

### Cell viability measurements

To calculate the percentage of viable cells in cultures containing MMS, the drug was added to exponentially growing *C. glabrata* or *S. cerevisiae* cultures at desired concentrations. At various time points thereafter, aliquots were removed, cells were counted using hemocytometer slides, and plated on drug-free YPD plates. Percentage viability was calculated based on the observed numbers of colonies relative to the corresponding cell counts.

### Fluorescent microscopy

The NLS-RFP construct was sub-cloned from plasmid pML85 (gift of Michael Lisby) into *C. glabrata* plasmid pMJ22 ^82^ (obtained from Addgene) using XhoI and NotI restriction sites. Slides for time-lapse microscopy were prepared by pipetting warm YPD containing 1% low melting agarose (with or without 0.03% MMS) onto glass slides and letting it solidify, forming YPD-agarose pads. Exponentially growing *C. glabrata* cells carrying the NLS-RFP plasmid were pipetted onto the YPD-agarose pads, covered with cover slips, and sealed using Biotium coverslip sealant (Fisher Scientific). The cells were imaged at room temperature for 6 or 10 hours at 10-minute intervals using a Nikon Eclipse Ti2 inverted microscope and Hamamatsu ORCA-Flash4.0 camera and analyzed using NIS-Elements software.

### Transcriptome analysis

Exponentially growing *C. glabrata* and *S. cerevisiae* cells were exposed to 0.1% MMS for one hour, at which point cells were harvested and total RNA was extracted using the RNeasy kit (Qiagen). The RNA samples were sent to Genewiz (https://www.genewiz.com) for RNAseq. Three biological replicates for each condition were submitted, but one “*S. cerevisiae* no MMS” sample failed quality control and was not processed further. The RNA-seq data were analyzed using Basepair software (https://www.basepairtech.com/) with a pipeline that included the following steps. Reads were aligned to the transcriptome derived from sacCer3 using STAR with default parameters. Read counts for each transcript was measured using featureCounts. Differentially expressed genes were determined using DESeq2 and a cut-off of 0.05 on adjusted p-value (corrected for multiple hypotheses testing) was used for creating differentially expressed gene lists. GSEA was performed on normalized gene expression counts, using gene permutations for calculating p-value. The list of *C. glabrata-S. cerevisiae* direct orthologs was downloaded from http://www.candidagenome.org/download/homology/orthologs and supplemented by manual curation of *C. glabrata* genes using http://www.candidagenome.org. Gene Ontology (GO) analysis was performed using FungiFun (https://elbe.hki-jena.de/fungifun) ^83^. Heatmaps were generated using R studio.

---

**Supplementary Figure 1.**
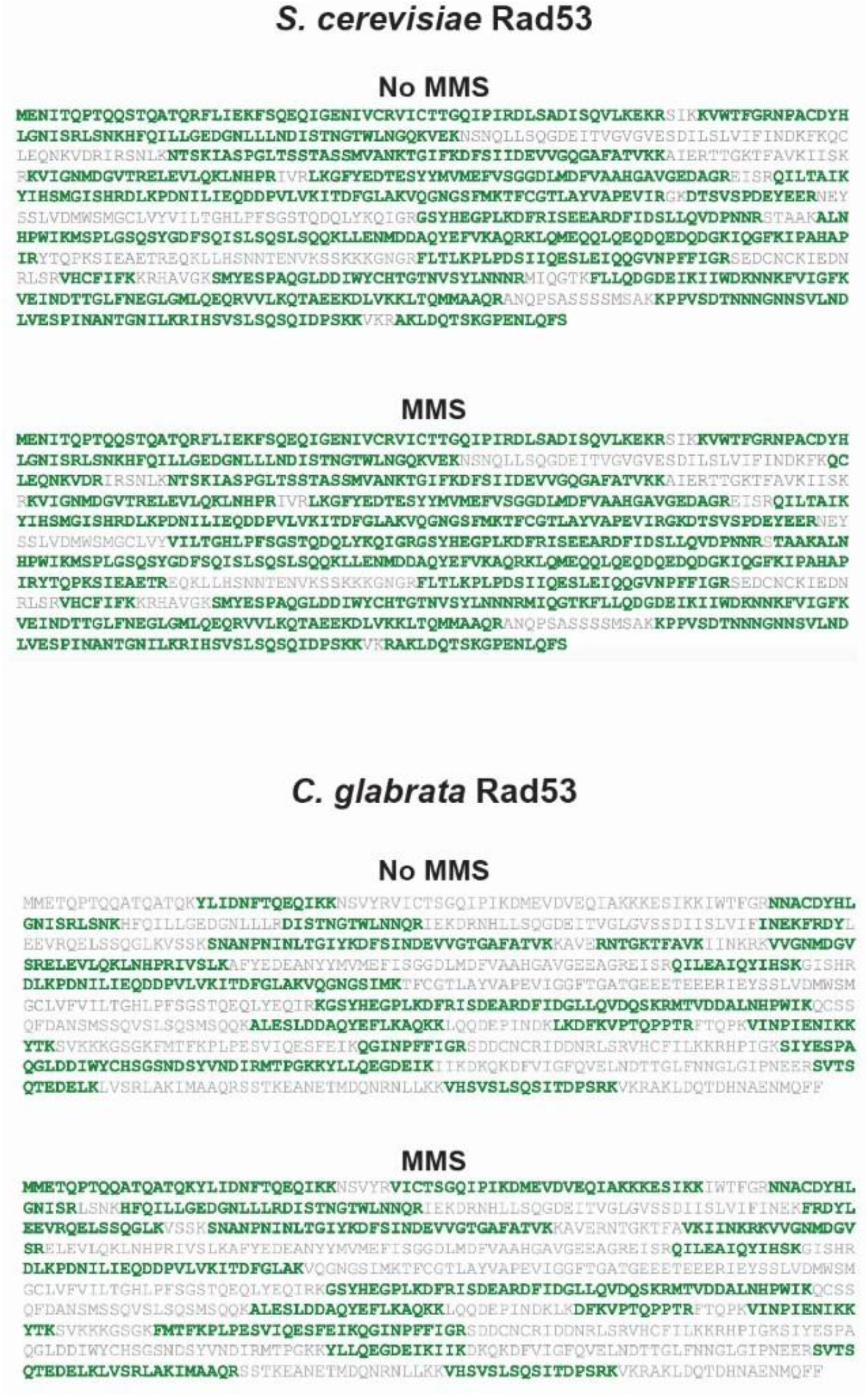
Peptides identified by MS in ScRad53 and CgRad53. (shown in green).

**Supplementary Figure 2.**
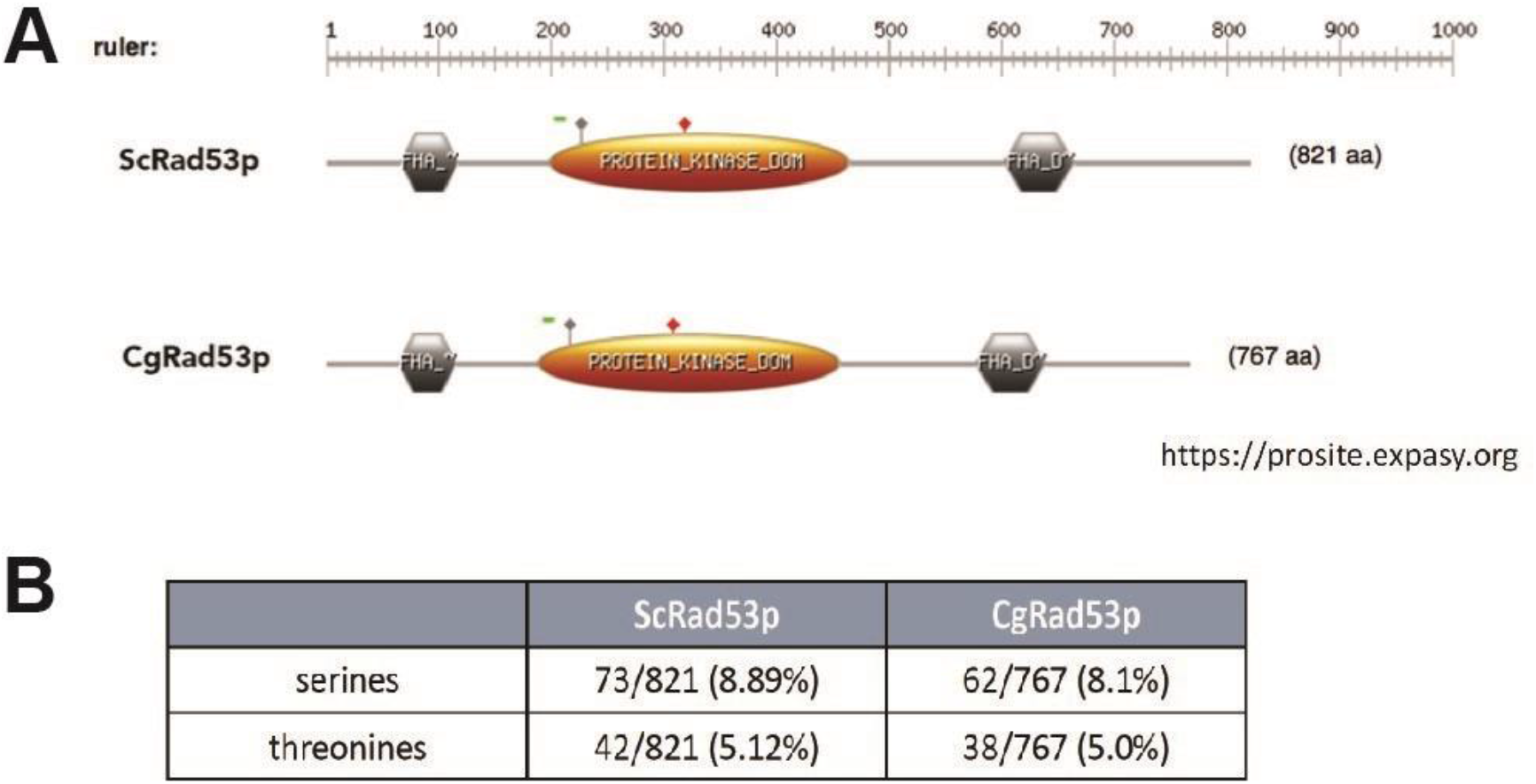
**a.** Overall Rad53 domain organization is similar in *S. cerevisiae* and *C. glabrata*. **b.** Overall percentage of serines and threonines is similar in ScRad53 and CgRad53.

**Supplementary Figure 3.**
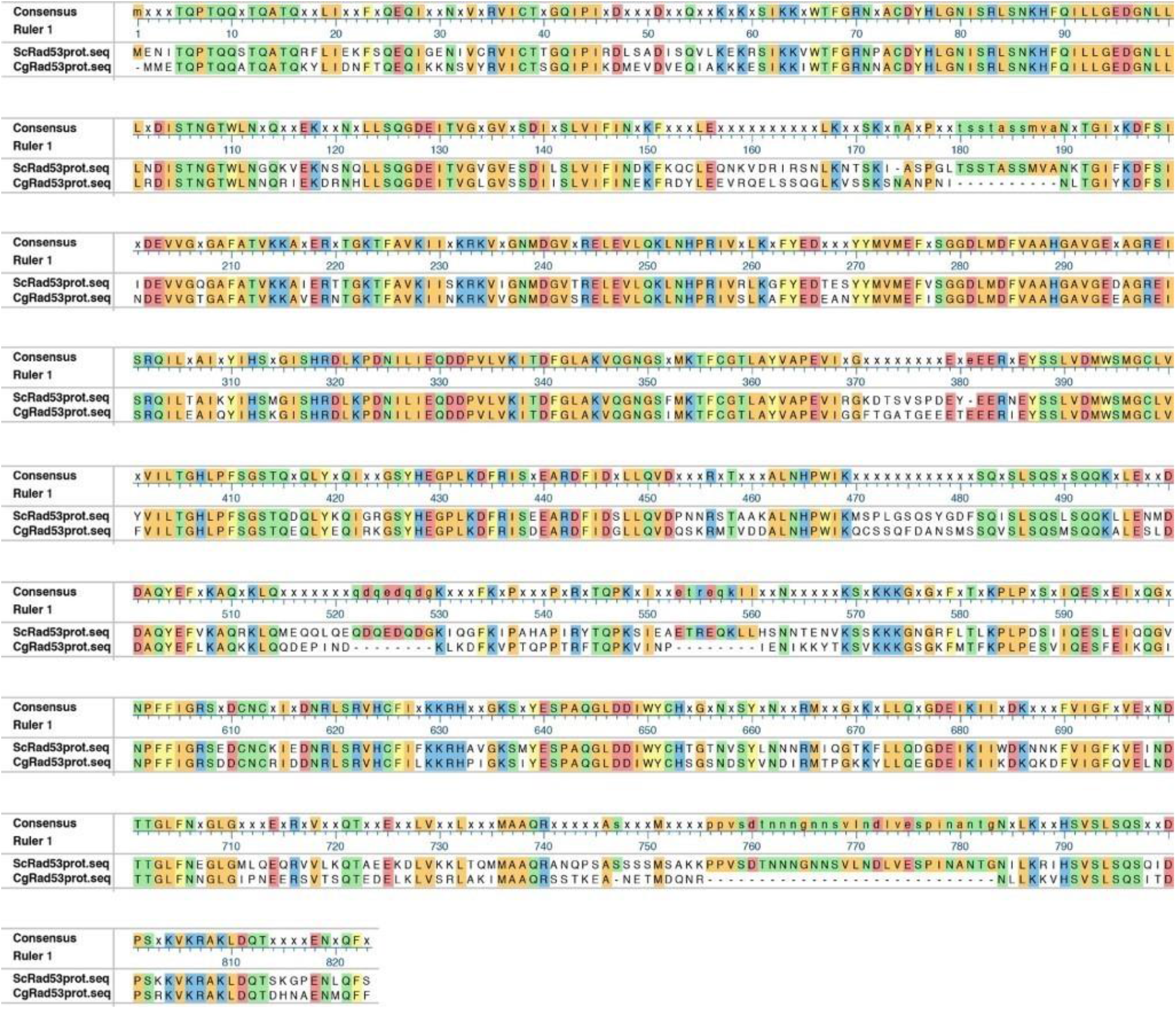
Rad53 alignment between *S. cerevisiae* and *C. glabrata* orthologs. The alignment was performed using MUSCLE in MegAlign Pro software (Lasergene).

**Supplementary Figure 4.**
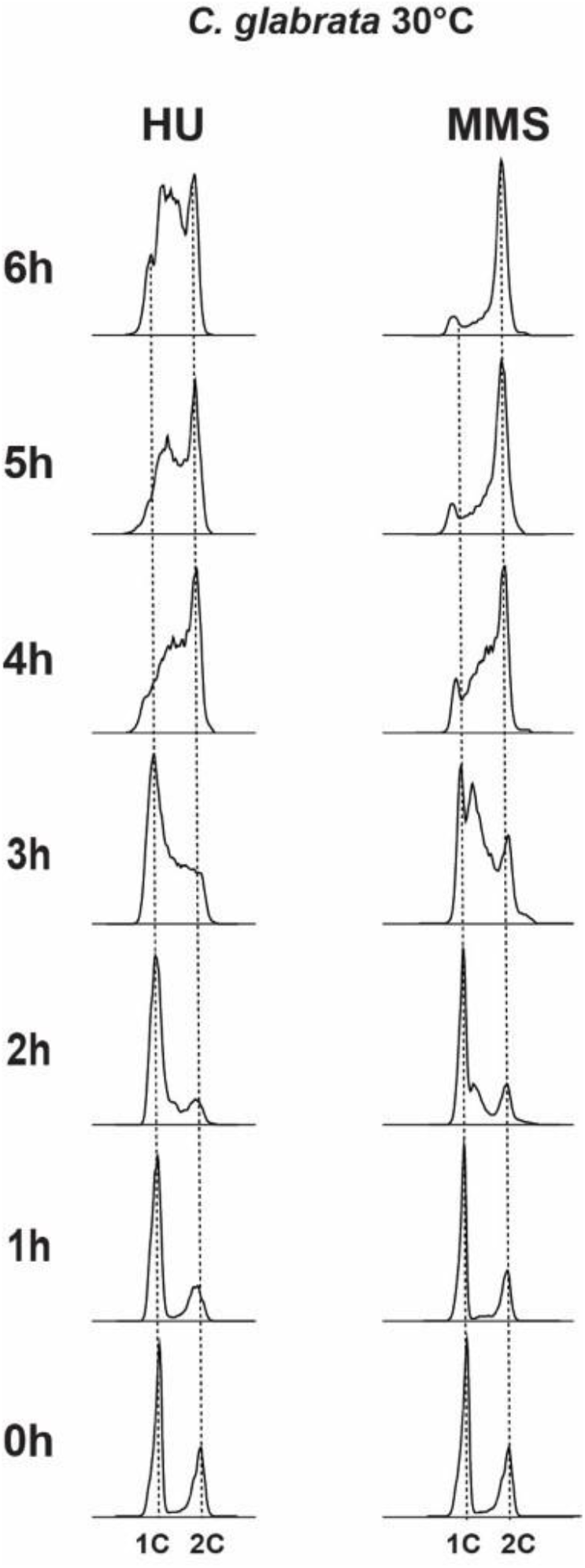
*C. glabrata* cells synchronized in G1 by carbon starvation and released into 0.03% MMS or 100 mM HU at 30°C behaved similarly to cells grown and released into these drugs at 37°C (Figure 4).

**Supplementary Figure 5.**
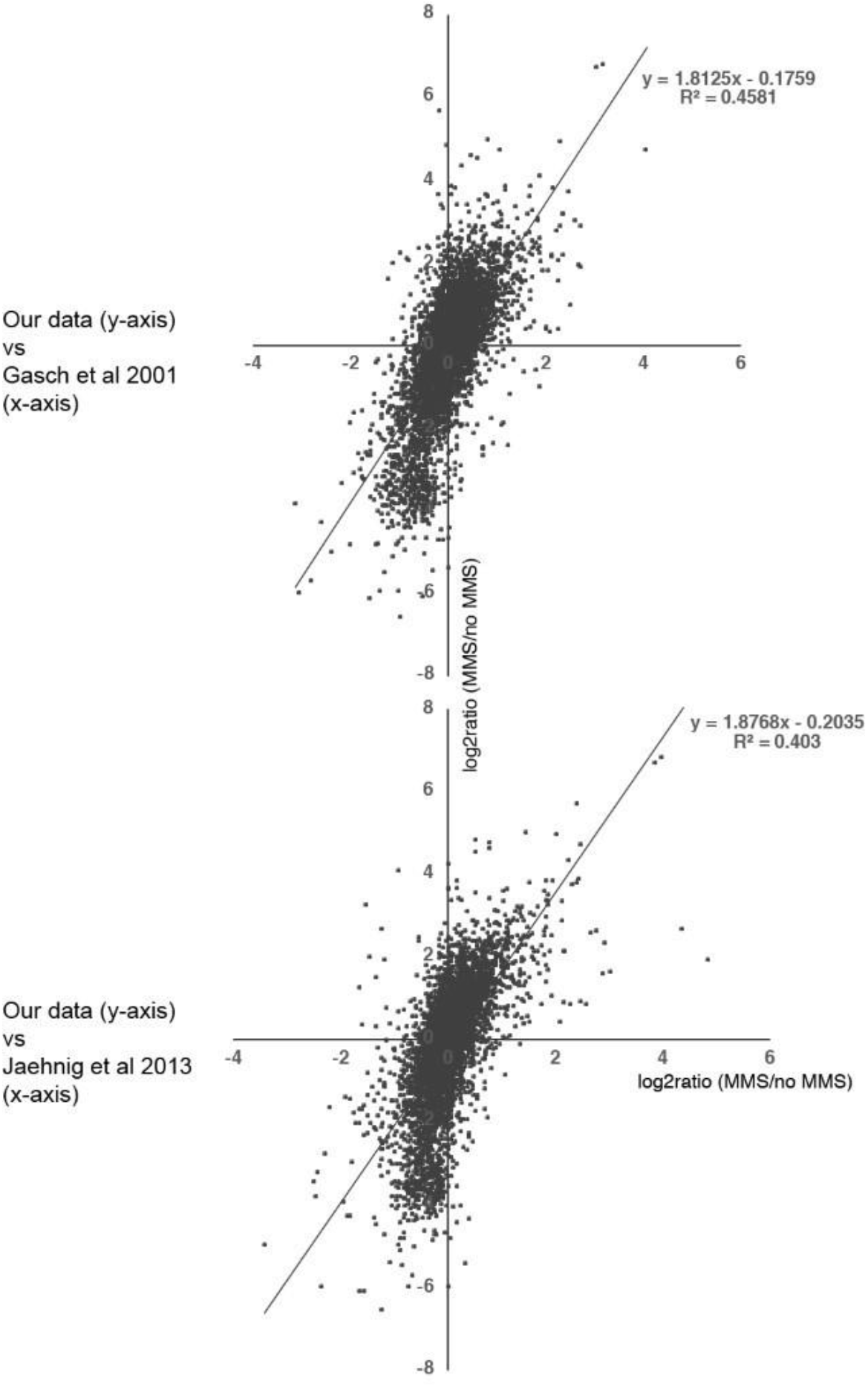
Good agreement between our transcriptomics data and those obtained by previous studies. The plots show correlations between our *S. cerevisiae* RNAseq data (log2ratio of MMS/noMMS for every gene) with those obtained by ^35^ and ^40^ The differences with our data are likely due to the fact that those studies used lower concentrations of MMS (0.02% or 0.03% vs 0.1% that we used) and because they were performed using microarray technology.

**Supplementary Figure 6.**
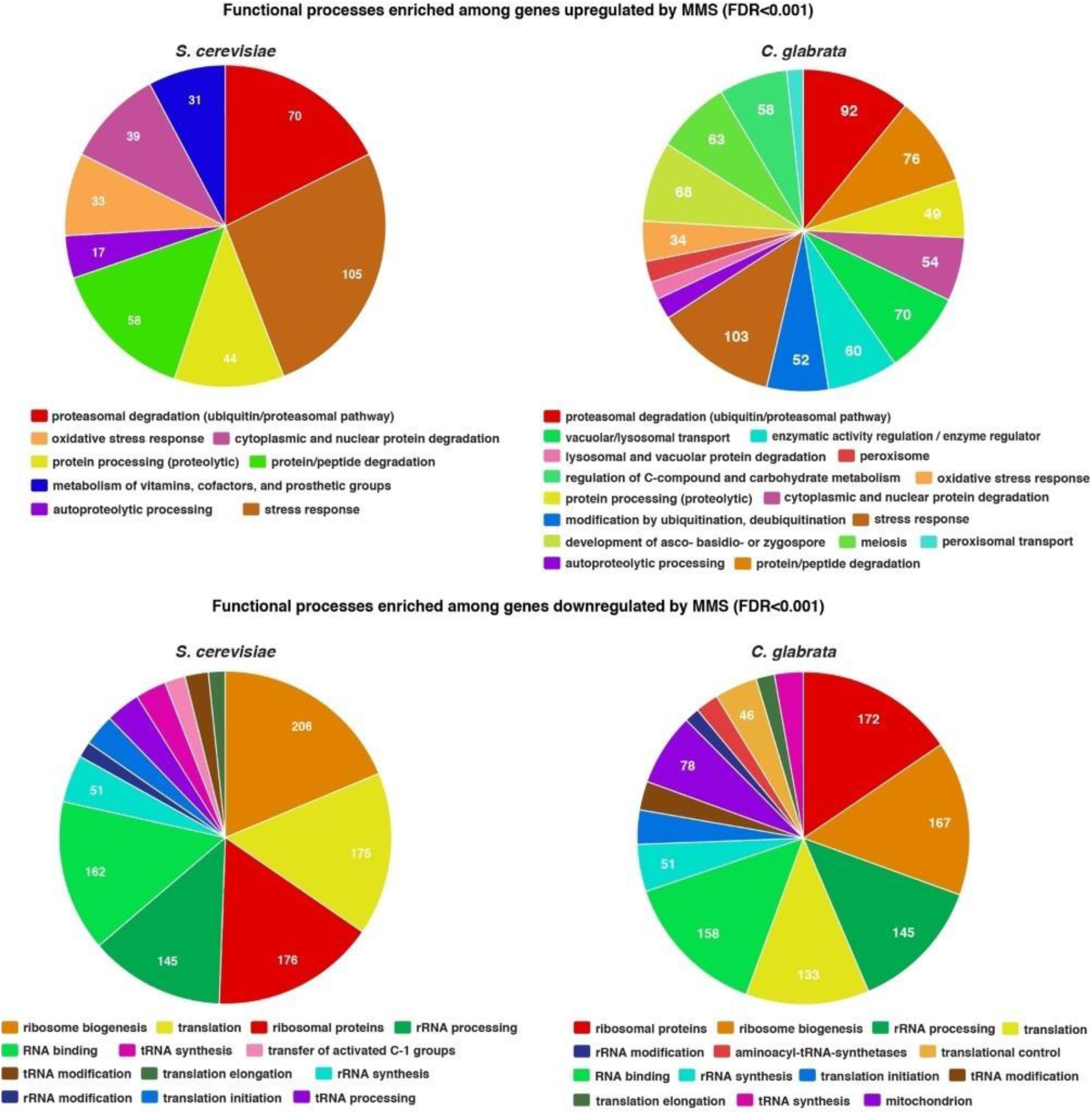
Gene Ontology (GO) analysis of genes upregulated and downregulated by MMS in *S. cerevisiae* and *C. glabrata*. Genes upregulated or downregulated at least two-fold in the presence of 0.1% MMS were subjected to GO analysis using the FungiFun tool (https://elbe.hki-jena.de/fungifun) ^83^.

**Supplementary Figure 7.**
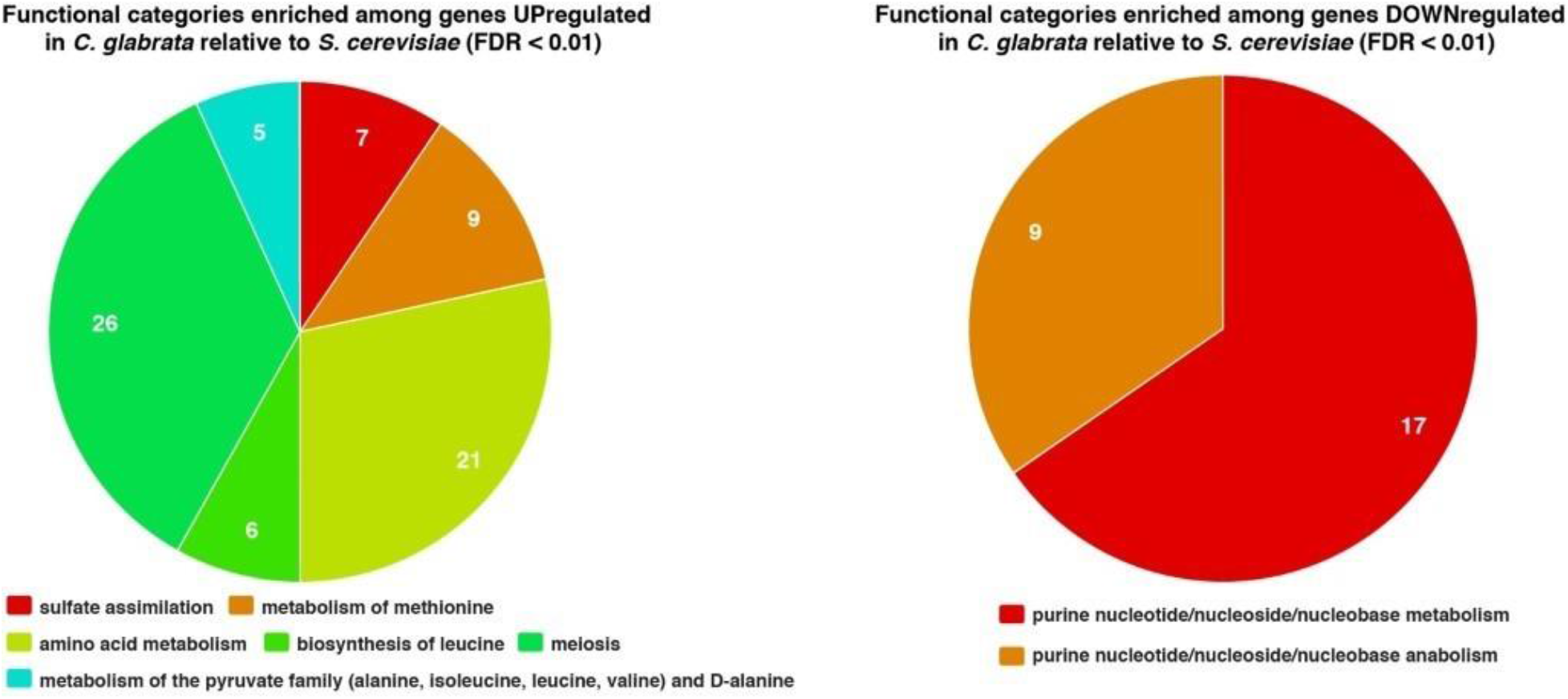
GO analysis of genes differentially regulated by MMS in *S. cerevisiae* and *C. glabrata*. Genes whose log2 ratio (MMS/noMMS) differed by at least 2 (reflecting at least a four-fold difference in the increase or decrease of transcript abundance in MMS) between *S. cerevisiae* and *C. glabrata* were subjected to GO analysis using the FungiFun tool (https://elbe.hki-jena.de/fungifun) ^83^.

## ACKNOWLEDGEMENTS

We thank Michael Lisby for the gift of plasmid pML85, the Rothstein lab for the gift of *S. cerevisiae* strain W4069-4C, Catherine Fox and Xiaolan Zhao for feedback on the manuscript, Marcella Lampon for technical assistance, and the Georgetown University Proteomics & Metabolomics Shared Resource for help with MS analysis. This work was supported by NIH 5R01AI109025 to DSP.

